# Electrophysiological signatures of acute systemic lipopolysaccharide: potential implications for delirium science

**DOI:** 10.1101/2020.11.25.398388

**Authors:** Ziyad W Sultan, Elizabeth R Jaeckel, Bryan M Krause, Sean M Grady, Caitlin A Murphy, Robert D Sanders, Matthew I Banks

**Author notes:** **Corresponding Author** Matthew I Banks, PhD, Professor, Department of Anesthesiology, 4605 Medical Sciences Center, 1300 University Avenue, Madison, WI 53706. these authors contributed equally (Ziyad W Sultan, Elizabeth R Jaeckel).

## Abstract

**Background:** Novel preventive therapies are needed for postoperative delirium, which especially affects aged patients. A mouse model is presented that captures inflammation-associated cortical slow wave activity (SWA) observed in patients, allowing exploration of the mechanistic role of prostaglandin-adenosine signaling.

**Methods:** EEG and cortical cytokine measurements (interleukin 6 [IL-6], monocyte chemoattractant protein-1 [MCP-1]) were obtained from adult and aged mice. Behavior, SWA, and functional connectivity (alpha-band weighted phase lag index) were assayed before and after systemic administration of lipopolysaccharide (LPS) +/- piroxicam (cyclooxygenase inhibitor) or caffeine (adenosine receptor antagonist). To avoid confounds from inflammation-driven changes in movement, which alter SWA and connectivity, electrophysiological recordings were classified as occurring during quiescence or movement, and propensity score matching used to match distributions of movement magnitude between baseline and LPS.

**Results:** LPS produces increases in cortical cytokines and behavioral quiescence. In movement-matched data, LPS produces increases in SWA (likelihood-ratio test: χ^2^(4)=21.51, p=0.00057), but not connectivity (χ^2^(4)=6.39, p=0.17). Increases in SWA associate with IL6 (p<0.001) and MCP-1 (p=0.001) and are suppressed by piroxicam (p<0.001) and caffeine (p=0.046). Aged animals compared to adult show similar LPS-induced SWA during movement, but exaggerated cytokine response and increased SWA during quiescence.

**Conclusions:** Cytokine-SWA correlations during wakefulness are consistent with observations in patients with delirium. Absence of connectivity effects after accounting for movement changes suggests decreased connectivity in patients is a biomarker of hypoactivity. Exaggerated effects in quiescent aged animals are consistent with increased hypoactive delirium in older patients. Prostaglandin-adenosine signaling may link inflammation to neural changes and hence delirium.

## Introduction

Inflammation is a key mechanism of many neurological disorders, be they chronic, such as dementia, or acute, such as delirium^1–6^. Even in less severe cases, acute inflammation affects brain function through illnesses including the common cold or influenza, causing neurological effects such as somnolence^7^. Elucidation of how inflammation affects brain function could highlight therapeutic targets to reduce the burden of these conditions.

Delirium is an acute disturbance of consciousness characterized by reduced attention, disorganized thinking, and fluctuating arousal levels that often affects sick elderly patients, especially those undergoing high risk surgery^8–12^. The electrophysiological hallmark of delirium is EEG slow wave activity (SWA)^13, 14^, similar to that observed during non-rapid eye movement sleep. SWA during delirium seems to particularly involve posterior brain regions^14^. Although evidence suggests that SWA during natural overnight sleep is restorative and enhances cognitive function, SWA during wakefulness, as occurs in delirium, is associated with cognitive deficits^15^. We have suggested that SWA may precipitate cognitive disintegration in delirium, such that patients are awake and confused^1^. Inflammation is the predominant acute cause of delirium^1, 14, 16^; it drives somnolence and enhances SWA in sleep^17–19^, and inflammation may similarly drive SWA in delirium^20^. In elderly patients, EEG SWA correlates with delirium severity, plasma cytokines, and EEG connectivity^14^. Our overarching hypothesis is that inflammation drives acute changes in SWA and disrupted cortical connectivity during wakefulness, triggering sudden and profound impairment in cognition. Herein, we test the link between inflammation and changes in brain activity and connectivity in a mouse model.

Progress on developing therapeutic interventions for delirium has been limited due to the lack of an established animal model to provide insights into its pathogenesis. The most critical limitation has been in identifying translational biomarkers of this complex human cognitive disorder. Recognizing the difficulty in establishing an animal model for these cognitive deficits, we focus on a translational biomarker of delirium, SWA in the EEG, building on previous work^21, 22^. In a critical advance from this earlier work, we focus on (i) SWA specific to active wakefulness, (ii) inflammation as the primary trigger, and (iii) how age may modulate these two factors, consistent with age being a key predisposing factor to delirium. Furthermore, we test interventions that attenuate the behavioral consequences of LPS. Based on prior studies showing that cyclooxygenase inhibitors attenuate acute behavioral changes induced by LPS by inhibiting the prostaglandin response^3, 23^, and that prostaglandins act as somnogens via adenosine signaling^24, 25^, we investigated the role of prostaglandin – adenosine signaling in linking inflammation to changes in neural activity and connectivity.

## Materials and Methods

Further methodological details can be found in Supplementary Methods online.

### Data collection

All procedures with animals were approved by the University of Wisconsin-Madison Institutional Animal Care and Use Committee (IACUC) and in full accordance with Research Animal Resources and Compliance (RARC). Adult (2-8 months old; n=68) and aged (16-24 months old, n=14) c57Bl/6J mice were used in this study (Supplementary Table 1). Of these 82 mice, 72 were instrumented for skull screw EEG recordings (bilateral parietal and frontal electrodes). After 5-7 days recovery, animal activity, resting-state EEG, and anterior-posterior functional connectivity were assayed during the animals’ dark (active) phase during a 1hr baseline period and for several hours after treatments, after which animals were euthanized and their brains frozen for later cytokine ELISA (Supplementary Figures 1 & 7). For LPS-alone experiments, animals were recorded 1 hour before and for 4 hours after intraperitoneal (IP) LPS administration at t=0hr (vehicle=0.9% NaCl, Low LPS=12.5 or 25µg/kg, High LPS=125µg/kg). For piroxicam experiments, recordings commenced at t=-2hr, piroxicam (10mg/kg IP) was administered at t=-1hr, followed by LPS (25µg/kg) at t=0hr, and euthanasia at t=4hr. Because caffeine has a short half-life in mice (<1hr)^26^, three IP injections of caffeine citrate (30mg/kg) were administered: the first at t=0hr (along with LPS 25µg/kg), then at t=1hr and t=2hr. To monitor movement and activity levels, video was recorded for the duration of electrophysiological recording and analyzed offline.

**Table 1.** Statistical modeling results. Shown are results of likelihood ratio tests for both the main models and models comparing caffeine experiments, along with associated sample counts (n_1_, n_2_), p-values and the results of pairwise comparisons. Effects of LPS were tested by comparing models with and without the group-by-epoch interaction (or group itself for cytokine data) using likelihood ratio tests. Caffeine experiments were fit in separate models together with equivalent saline controls; since there are only two factor levels no pairwise comparisons are needed. Post-hoc comparisons used the Kenward-Roger method and p-values for pairwise comparisons were adjusted for multiple comparisons by estimating a multivariate t-distribution using the emmeans package for R^36^.

| Measure | Likelihood Ratio Test /<br>Pairwise Comparison | n<sub>1</sub>,n<sub>2</sub> | p-value | Measure | Likelihood Ratio Test /<br>Pairwise Comparison | n<sub>1</sub>,n<sub>2</sub> | p-value |
| --- | --- | --- | --- | --- | --- | --- | --- |
| **A** IL-6 | $\chi^2(4) = 29.0$ | | <0.0001 | **B** MCP-1 | $\chi^2(4) = 29.1$ | | <0.0001 |
|  | VEH vs Low LPS | 5,18 | 0.68 |  | VEH vs Low LPS | 5,18 | 0.87 |
|  | VEH vs High LPS | 5,8 | 0.0004 |  | VEH vs High LPS | 5,8 | 0.0003 |
|  | Low LPS vs High LPS | 18,8 | 0.0002 |  | Low LPS vs High LPS | 18,8 | <0.0001 |
|  | VEH vs PXM+Low LPS | 5,12 | 0.23 |  | VEH vs PXM+Low LPS | 5,12 | 0.12 |
|  | Low LPS vs PXM+Low LPS | 18,12 | 0.69 |  | Low LPS vs PXM+Low LPS | 18,12 | 0.17 |
|  | Low LPS vs Aged Low LPS | 18,13 | 0.0033 |  | Low LPS vs Aged Low LPS | 18,13 | 0.014 |
| **C** SWA | $\chi^2(4) = 87.8$ | | <0.0001 | **D** gamma | $\chi^2(4) = 78.6$ | | <0.0001 |
|  | VEH vs Low LPS | 7,17 | <0.0001 |  | VEH vs Low LPS | 7,17 | <0.0001 |
|  | VEH vs High LPS | 7,8 | <0.0001 |  | VEH vs High LPS | 7,8 | <0.0001 |
|  | Low LPS vs High LPS | 17,8 | 0.041 |  | Low LPS vs High LPS | 17,8 | 0.043 |
|  | VEH vs PXM+Low LPS | 7,8 | 0.72 |  | VEH vs PXM+Low LPS | 7,8 | 0.5 |
|  | Low LPS vs PXM+Low LPS | 17,8 | <0.0001 |  | Low LPS vs PXM+Low LPS | 17,8 | <0.0001 |
|  | Low LPS vs Aged Low LPS | 17,14 | 0.025 |  | Low LPS vs Aged Low LPS | 17,14 | 0.36 |
| Caffeine | $\chi^2(1) = 36.6$ | 8,10 | <0.001 | Caffeine | $\chi^2(1) = 61.5$ | 8,10 | <0.0001 |
| **E** wPLI | $\chi^2(4) = 26.2$ | | <0.0001 | **F** Movt | $\chi^2(4) = 18.8$ | | 0.00085 |
|  | VEH vs Low LPS | 7,17 | 0.011 |  | VEH vs Low LPS | 7,17 | 0.014 |
|  | VEH vs High LPS | 7,8 | 0.002 |  | VEH vs High LPS | 7,8 | 0.0005 |
|  | Low LPS vs High LPS | 17,8 | 0.61 |  | Low LPS vs High LPS | 17,8 | 0.25 |
|  | VEH vs PXM+Low LPS | 7,8 | 0.063 |  | VEH vs PXM+Low LPS | 7,8 | 0.23 |
|  | Low LPS vs PXM+Low LPS | 17,8 | 1 |  | Low LPS vs PXM+Low LPS | 17,8 | 0.83 |
|  | Low LPS vs Aged Low LPS | 17,12 | 0.031 |  | Low LPS vs Aged Low LPS | 17,14 | 0.99 |
| Caffeine | $\chi^2(1) = 18.2$ | 8,10 | 0.0002 | Caffeine | $\chi^2(1) = 18.8$ | 8,10 | <0.0001 |
| **G** SWA Movt-matched | $\chi^2(4) = 21.5$ | | 0.00025 | **H** SWA Quiescent | $\chi^2(4) = 19.7$ | | 0.00057 |
|  | VEH vs Low LPS | 7,16 | 0.0086 |  | VEH vs Low LPS | 6,16 | 0.67 |
|  | VEH vs High LPS | 7,5 | 0.0057 |  | VEH vs High LPS | 6,5 | 0.32 |
|  | Low LPS vs High LPS | 16,5 | 0.77 |  | Low LPS vs High LPS | 16,5 | 0.82 |
|  | VEH vs PXM+Low LPS | 7,7 | 1 |  | VEH vs PXM+Low LPS | 6,7 | 1 |
|  | Low LPS vs PXM+Low LPS | 16,7 | 0.013 |  | Low LPS vs PXM+Low LPS | 16,7 | 0.83 |
|  | Low LPS vs Aged Low LPS | 16,13 | 0.99 |  | Low LPS vs Aged Low LPS | 16,13 | 0.0086 |
| Caffeine | $\chi^2(1) = 3.98$ | 8,8 | 0.046 | Caffeine | $\chi^2(1) = 2.61$ | 8,8 | 0.11 |
| **I** wPLI Movt-matched | $\chi^2(4) = 6.39$ | | 0.17 | **J** wPLI Quiescent | $\chi^2(4) = 4.77$ | | 0.31 |
| Caffeine | $\chi^2(1) = 7.0$ | 8,6 | 0.0082 | Caffeine | $\chi^2(1) = 1.49$ | 8,8 | 0.22 |

**Figure 1.**
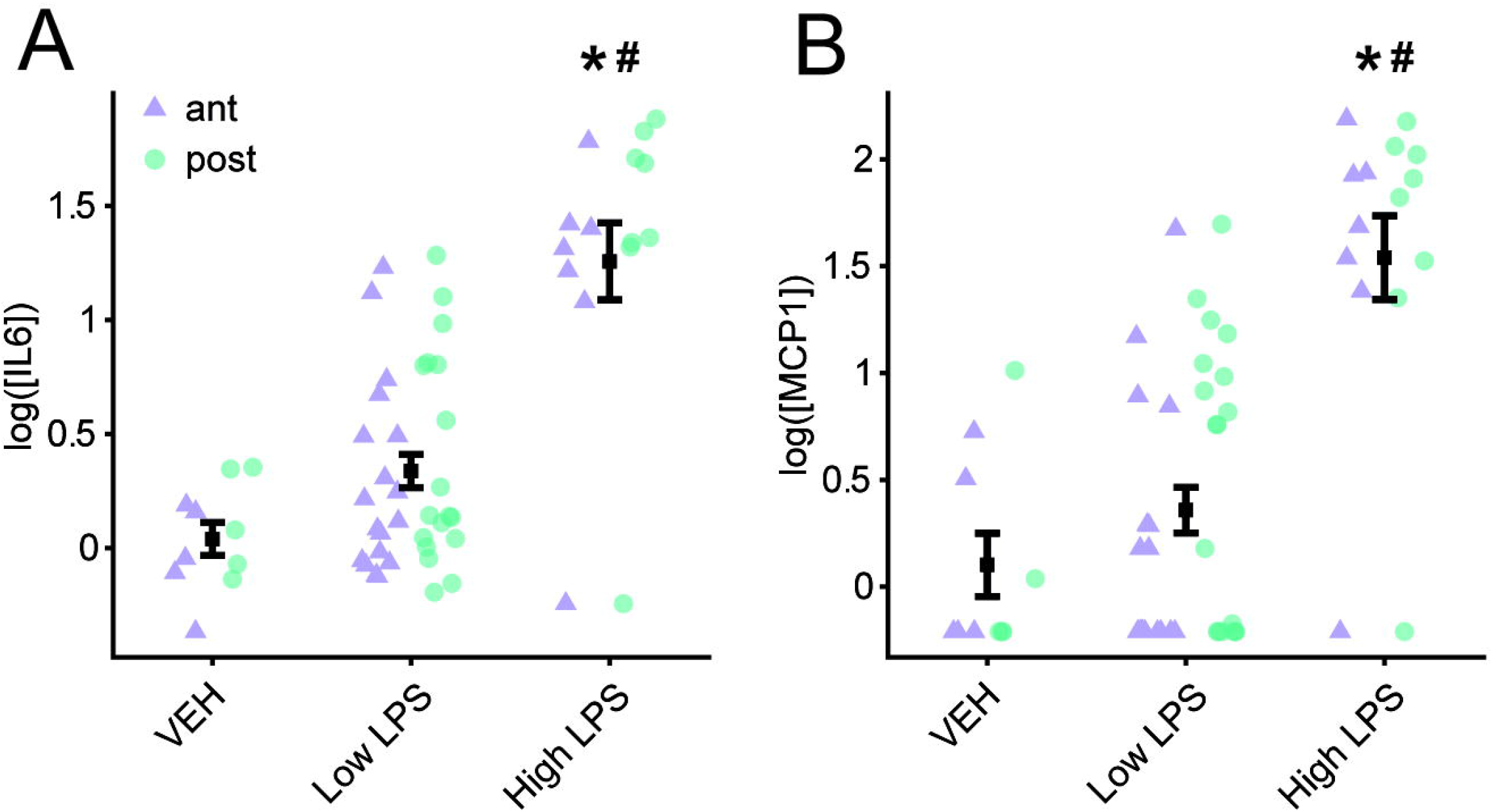
Proinflammatory cytokine levels in neocortex after LPS treatment. **A**. Shown are interleukin-6 (IL-6) protein concentrations measured via ELISA in neocortical homogenate samples from mice (including animals with and without EEG implant) euthanized four hours after LPS injection. Each point represents log IL-6 concentration (pg/ml) from samples obtained bilaterally from anterior (ant; *purple triangles*) or posterior cortex (post; *green circles*). Overlaid symbols (*black*) represent the within-group mean across all samples. Error bars represent ± SEM. * indicates significant difference from Vehicle. # indicates significant difference from Low LPS. **B**. Monocyte chemoattractant protein-1 (MCP-1) protein concentration values are shown. MCP-1 samples below the threshold of detection were set to the square root of the lowest observed quantity (1.13). LPS, lipopolysaccharide; VEH, vehicle.

### Data analysis

Band power analysis of EEG data proceeded according to standard techniques^27^, with power calculated in 4-second sliding windows in the delta (i.e. SWA, 2-4Hz), theta (4-12Hz), alpha (13-20Hz), beta (20-30Hz), and gamma (30-80Hz) bands and normalized by total power. Functional connectivity was assayed using the alpha band debiased weighted phase lag index (wPLI)^28^, calculated in 20-second sliding windows between anterior and posterior channel pairs for each hemisphere, and averaged across hemispheres. We chose alpha band wPLI *a priori* because it is a standard metric of functional connectivity^27^ and is used especially in other papers on delirium^14, 29, 30^. A movement signal, derived from the smoothed and normalized frame-by-frame video difference signal, was aligned in time with the simultaneously recorded EEG signal, and the movement signal averaged in each 4-sec or 20-sec epoch for band power and wPLI analysis, respectively. Epochs with nonzero estimated movement were used to calculate electrophysiological parameters corresponding to active wakefulness. To ensure that drug-induced changes in activity level for epochs classified as active wakefulness did not influence measured electrophysiological parameters, distributions of movement signal magnitude were matched between baseline and treatment periods using propensity score matching (PSM; Supplementary Figure 2)^31^.

**Figure 2.**
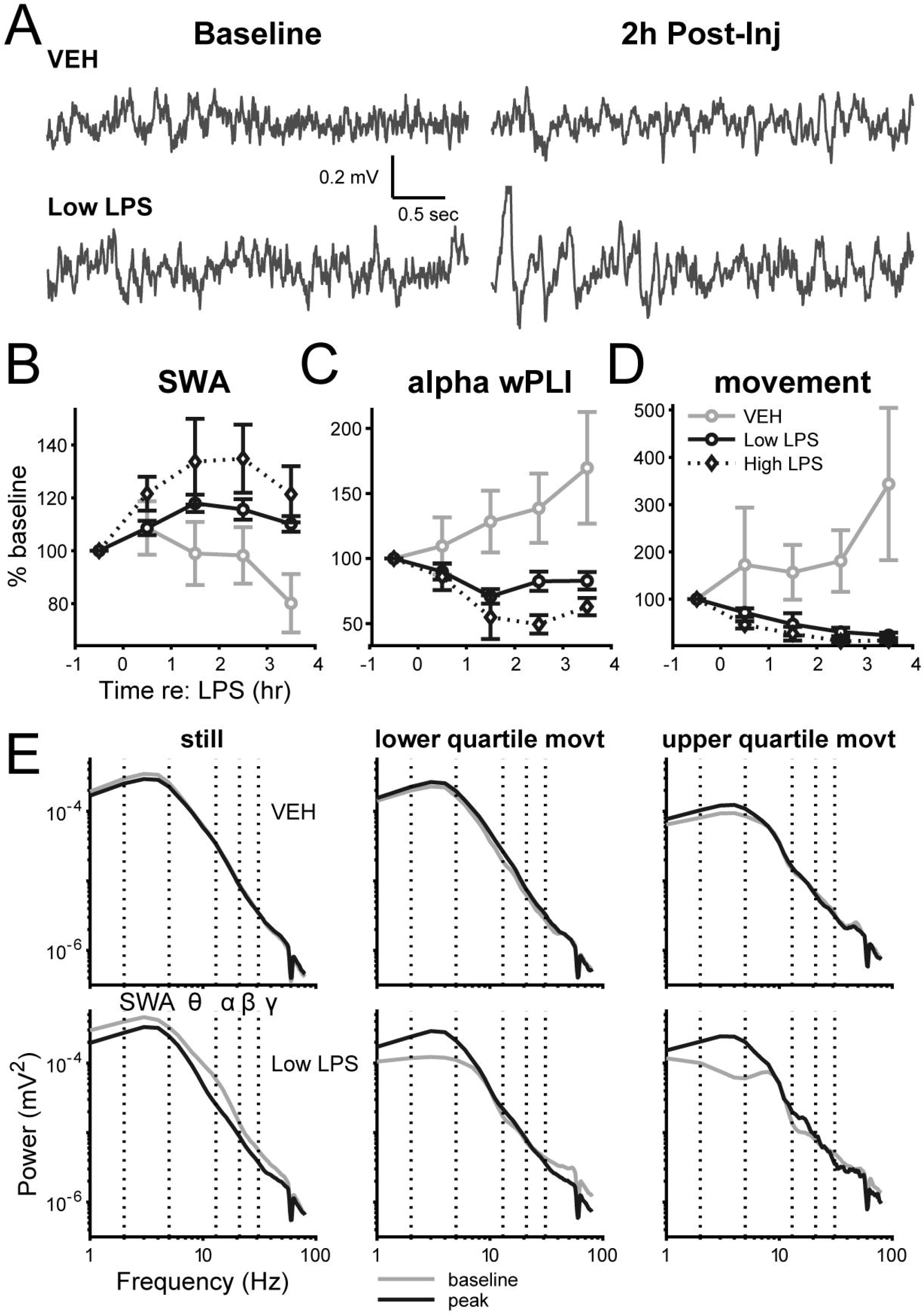
Generalized slowing of EEG and disrupted functional connectivity following LPS injection. **A**. Representative time-domain EEG signals during the pre-injection baseline hour (*left*) and two hours post-injection (*right*) are shown for animals that received either a saline “Vehicle” (*top*) or 25 µg/kg “Low” LPS injection (*bottom*). Traces were selected based on having mean SWA values approximately equal to the mean SWA over the entire hour. **B**. The time series of SWA (2-4Hz) normalized to mean spectral power (2-80Hz) are shown for the different LPS doses. Symbols represent the mean percent change in SWA from baseline across all animals at each LPS dose at each recording hour. Error bars represent ± SEM. **C**. Time series of alpha-band (13-20Hz) anterior-posterior weighted-phase lag index (wPLI), a measure of functional connectivity. **D**. The time series of movement for each LPS dose is shown. Symbols represent the mean percent change in movement from baseline across all animals at each dose of LPS at each recording hour. Error bars represent ± SEM. **E**. Example EEG power spectra separated according to movement magnitude. Power spectra were calculated in overlapping 4-second windows and aligned with movement epochs, then data were binned into quiescence (i.e. zero movement, *left*), lower quartile movement (*middle*), or upper quartile movement (*right*), and averaged. Pre-injection spectra (“baseline”; *gray*) and average spectra of 1-3 hours post-injection (“peak”; *black*), averaged across four EEG channels, are shown from animals that received either Vehicle (*top*) or a 25 µg/kg “Low” dose of LPS (*bottom*). LPS, lipopolysaccharide; SWA, slow-wave activity; VEH, vehicle; wPLI, weighted phase lag index.

Cytokine quantification was applied to brains from 46 mice with EEG recordings and 10 uninstrumented mice subjected to identical drug treatments (Supplementary Table 1). The cytokines interleukin 6 (IL-6) and monocyte chemoattractant protein-1 (MCP-1) were quantified by multiplex ELISA performed by Eve Technologies (#MDF10, Calgary, AB, Canada). IL-6 was selected *a priori* as the primary cytokine of interest, as brain levels of IL-6 increase dramatically in the hours following peripheral treatment with LPS^32^, with IL-6 having a relatively long half-life compared to other cytokines^33^, and IL-6 tends to be higher in hip-fracture surgical patients with delirium compared to those without^34^. Additionally, MCP-1 was chosen based on our recent findings in humans showing that MCP-1 correlates with delirium severity and EEG slow wave activity^14^.

To compare LPS-driven changes in EEG and activity measures across time, data were averaged across epochs during a “peak effect” period (t=1 to 3hrs post-injection) and compared to the average across epochs during the “baseline” pre-injection period (t=-1 to 0hr). LPS effects on band power, movement, and wPLI were assessed by fitting linear mixed effects models^35^. Fixed effects were treatment group, experiment epoch (baseline vs. peak effect), and their interaction, with random effects for animal to account for repeated measures and for multiple electrodes within animals when appropriate. Models fit to cytokine data used group and region (anterior/posterior) as fixed effects.

LPS dose categories were as follows: “Vehicle” (n=7) corresponded to 0µg/kg (i.e. 0.9% NaCl alone), “Low LPS” to 12.5µg/kg (n=8) and 25µg/kg (n=15) (grouped together as the outcomes of statistical analyses were unchanged by grouping the low doses), and “High LPS” (n=8) to 125µg/kg. Effects of LPS were tested by comparing models with and without the group-by-epoch interaction (or group-by-region for cytokine data) using likelihood ratio tests. Caffeine and the associated saline controls were fit in separate models from the other groups due to their differing experiment schedule, which may have affected behavior and precluded direct comparisons of these groups to the others. Post-hoc comparisons used the Kenward-Roger method and p-values were adjusted for the family of relevant multiple comparisons by estimating a multivariate t-distribution using the emmeans package for R^36^.

For relationships between cytokine levels and changes in movement-matched SWA, we fit a linear model to all data in the Vehicle, Low LPS, and High LPS groups to estimate SWA as a function of cytokine concentration. We then predicted SWA changes based on cytokine levels observed in the PXM + Low LPS group and tested whether the mean residual differed from zero using a one-sample t-test.

## Results

### LPS injection increases inflammatory markers in the brain

We first used ELISA to measure protein levels of proinflammatory cytokines IL-6 and MCP-1 (Figure 1A-B) in anterior or posterior mouse neocortex (*purple* and *green*, respectively) four hours after IP injection of LPS. There was no interaction between region and LPS dose for IL-6 (likelihood ratio test adding location: χ^2^(4)=4.30, p=0.37) or MCP-1 (likelihood ratio test adding location: χ^2^(4)=6.85, p=0.14); thus, we averaged anterior and posterior samples. IL-6 and MCP-1 levels were elevated following LPS injection, and there was a significant overall effect of LPS group on cytokine concentration (Table 1A-B). Cytokine levels in Vehicle animals were not different from Low LPS animals, though both groups had significantly lower cytokine levels compared to High LPS.

### LPS injection increases SWA and decreases antero-posterior connectivity

LPS administration was followed by a slowing of resting-state brain activity, manifest as an increase in SWA in the EEG signal (Figure 2A-B) and a concomitant decrease in amplitude in higher frequency bands, such as gamma (Supplementary Figure 3). SWA band power showed clear changes following LPS injection, with the effect reaching a peak between 1- and 3-hours post-injection (“peak LPS”; Figure 2B). Increases in SWA from baseline to peak LPS significantly depended on LPS dose (Table 1C). Similarly, decreases in gamma power following LPS treatment were dose-dependent (Supplementary Figure 4A; Table 1D). In a regional analysis of SWA we found a significant interaction between anterior/posterior region, group, and time (likelihood ratio test: χ^2^(3)=8.82, p=0.032). However, this effect was limited to a larger posterior compared to anterior increase in the High LPS animals, which could be explained by ceiling effects specifically in the High LPS condition (at baseline, anterior SWA was greater than posterior power in all groups). Since animals in all subsequent experiments received either Vehicle or Low LPS, and because the LPS effect was comparable anterior and posterior in the Low LPS group, we did not alter our *a priori* statistical plan to combine anterior and posterior channels in analyses of power.

**Figure 3.**
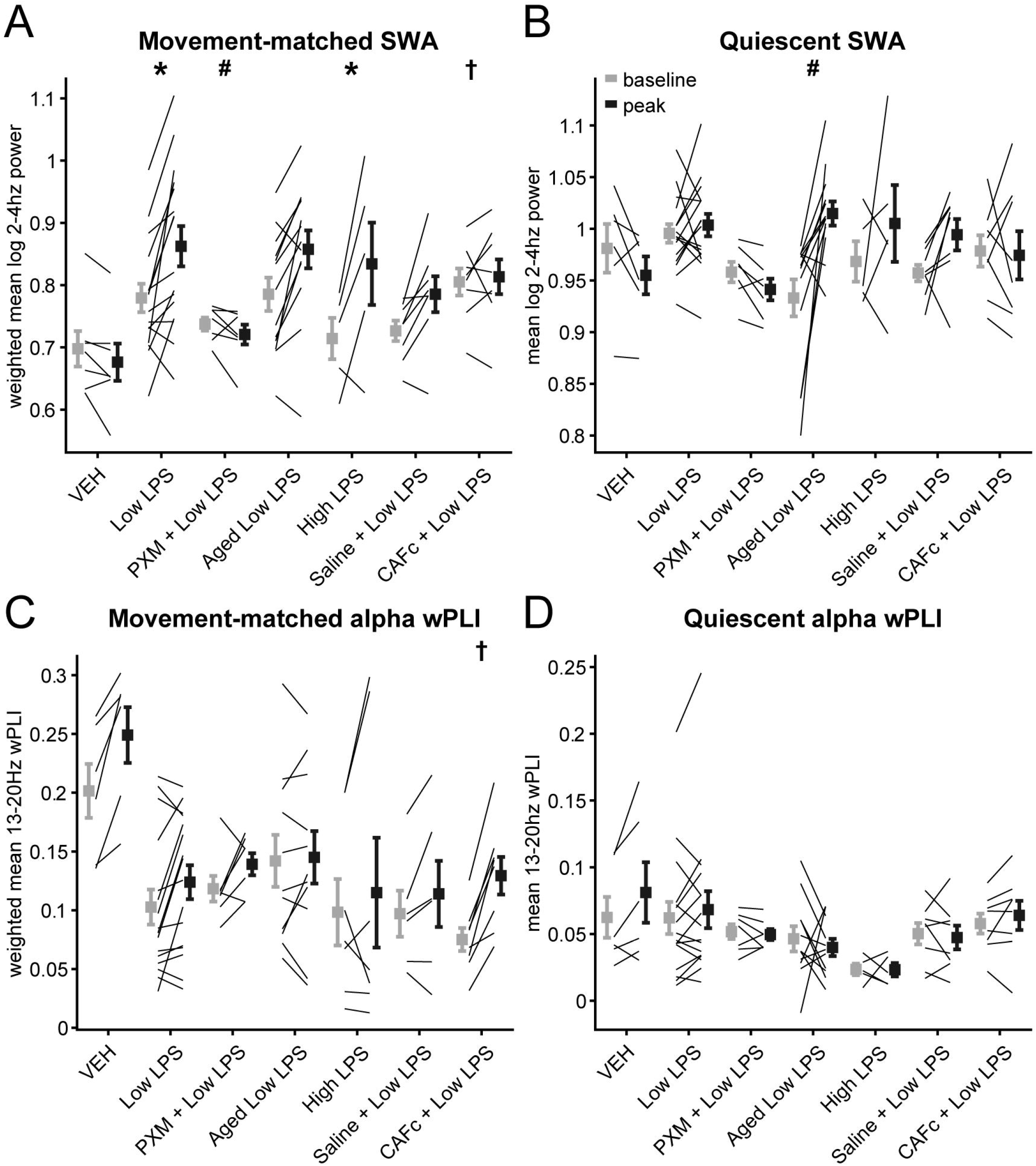
Changes in SWA and alpha-band wPLI during movement or quiescence. **A**. Movement-matched SWA values at baseline (*gray*) or peak LPS (*black*). Lines indicate mean log SWA values for individual animals while symbols represent group averages. Error bars indicate ± SEM. * indicates significant difference of differences from Vehicle. # indicates significant difference of differences from Low LPS. † indicates significant difference of differences from Saline + Low LPS. **B**. SWA during quiescence. **C**. Movement-matched alpha-band wPLI. **D**. Alpha wPLI during quiescence. CAFc, caffeine citrate; LPS, lipopolysaccharide; PXM, piroxicam; SWA, slow-wave activity; VEH, vehicle; wPLI, weighted phase lag index.

**Figure 4.**
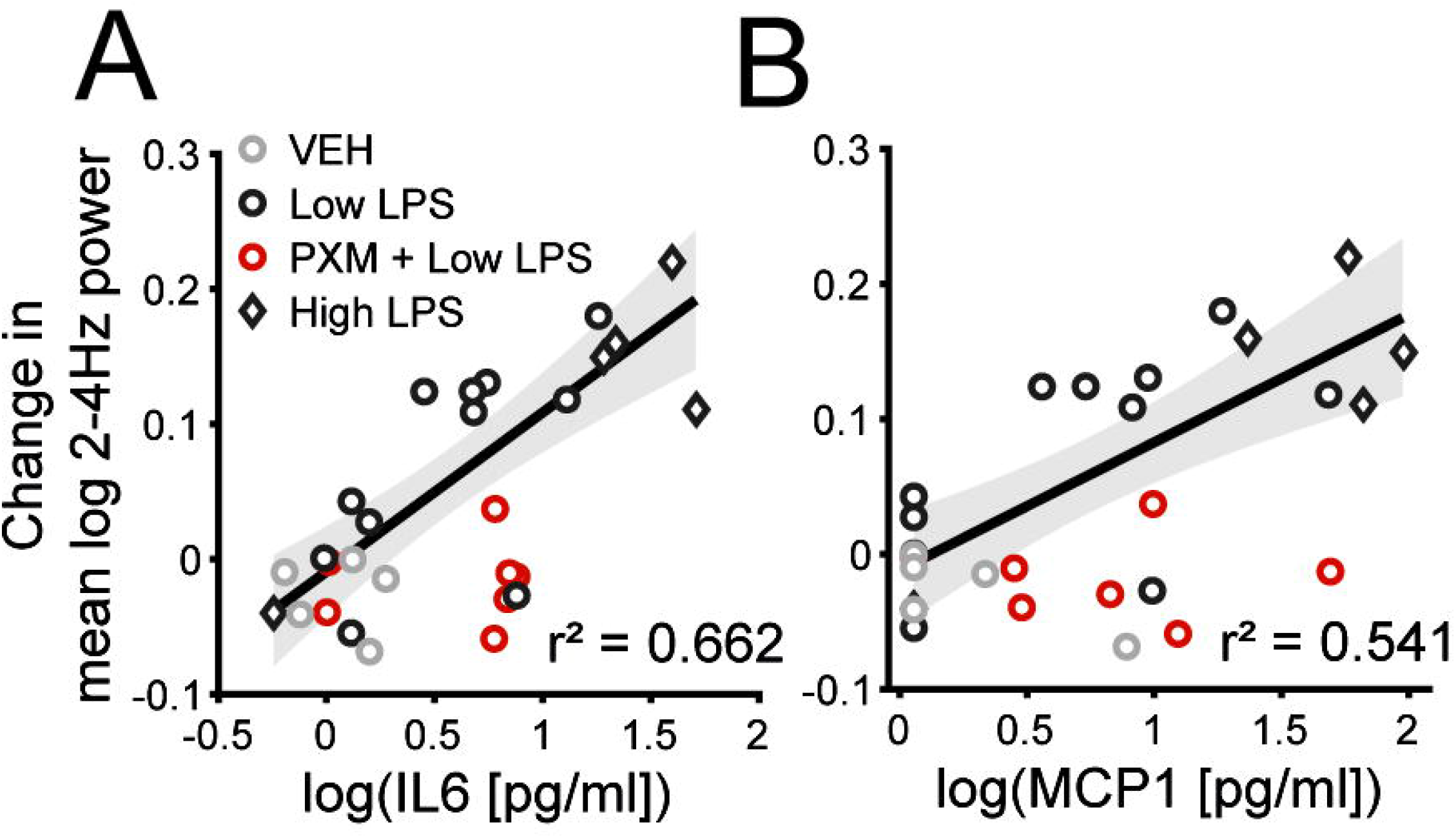
Cytokine correlations with changes in movement-matched SWA. **A**. Scatterplot of the change in movement matched SWA from baseline to peak LPS versus log IL-6 concentration. Markers represent values for individual animals. The line indicates the polynomial least-squares fit (using the MATLAB function “regress”) for LPS-only groups, and the shading indicates the 95% prediction interval of the regression line. The R^2^ value for the fit is also indicated. **B**. Scatterplot of the change in movement matched SWA from baseline to peak versus log MCP-1 concentration. IL-6, interleukin-6; LPS, lipopolysaccharide; MCP-1, Monocyte chemoattractant protein-1; PXM, piroxicam; SWA, slow-wave activity; VEH, vehicle.

Previous reports have suggested that cortical functional connectivity (measured by alpha-band wPLI) is disrupted in delirious patients^14, 29^, and systemic LPS can alter connectivity in human volunteers^37^. Consistent with these previous observations, we observed decreased alpha-band wPLI following injection of LPS (Figure 2C). Alpha-band connectivity decreased more in Low LPS and High LPS animals compared to Vehicle, though there was no difference between LPS doses (Table 1E).

### LPS decreases movement

Animals injected with LPS exhibited sickness behavior typical of systemic inflammation, including piloerection, hunched posture, and reduced locomotion and grooming activity^38^. The effect of LPS on overall activity level was quantified by the magnitude of the movement signal derived from the video recordings (Supplementary Figure 5, *black*; see Methods). Consistent with prior observations^39^, LPS caused a dramatic decrease in movement from baseline to peak LPS hours (Figure 2D). Movement decreased more in Low LPS and High LPS animals compared to Vehicle, though there was no significant difference between LPS doses (Table 1F).

**Figure 5.**
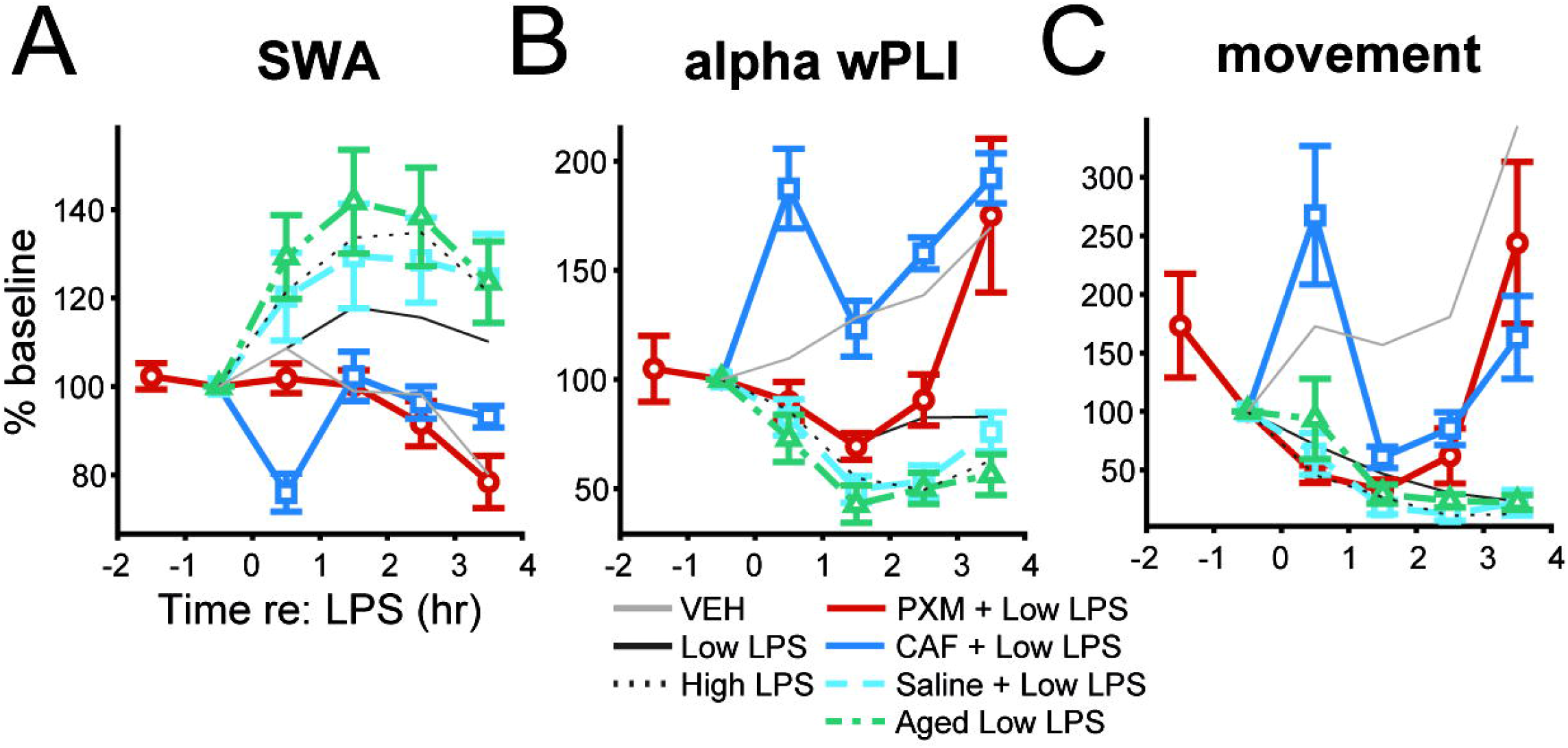
EEG and movement time course summary across groups. **A**. The time series of SWA (2-4Hz) normalized to mean spectral power (2-80Hz) are shown for all LPS groups (Vehicle, Low LPS, and High LPS are the same as in Figure. 2). Symbols represent the mean percent change in SWA from baseline across all animals at each LPS dose at each recording hour. Error bars represent ± SEM. **B**. Gamma power time series. **C**. Alpha wPLI time series. **D**. Movement time series. Bars indicate ± SEM. CAFc, caffeine citrate; LPS, lipopolysaccharide; PXM, piroxicam; SWA, slow-wave activity; VEH, vehicle; wPLI, weighted phase lag index.

### SWA in LPS increases after correcting for changes in movement

Because EEG delta power is higher and gamma power is lower during sleep and quiescent wakefulness compared to active wakefulness^40, 41^, the changes in movement following LPS injection could themselves account for the observed slowing in EEG signals. In all animals, movement was negatively correlated with SWA (Supplementary Figure 5; Supplementary Figure 6A; r^2=^0.313, p<0.05). Because delirium in patients is characterized by elevated delta power during wakefulness^13, 14^, we tested whether LPS caused a slowing of the EEG after accounting for the LPS-induced decrease in movement.

**Figure 6.**
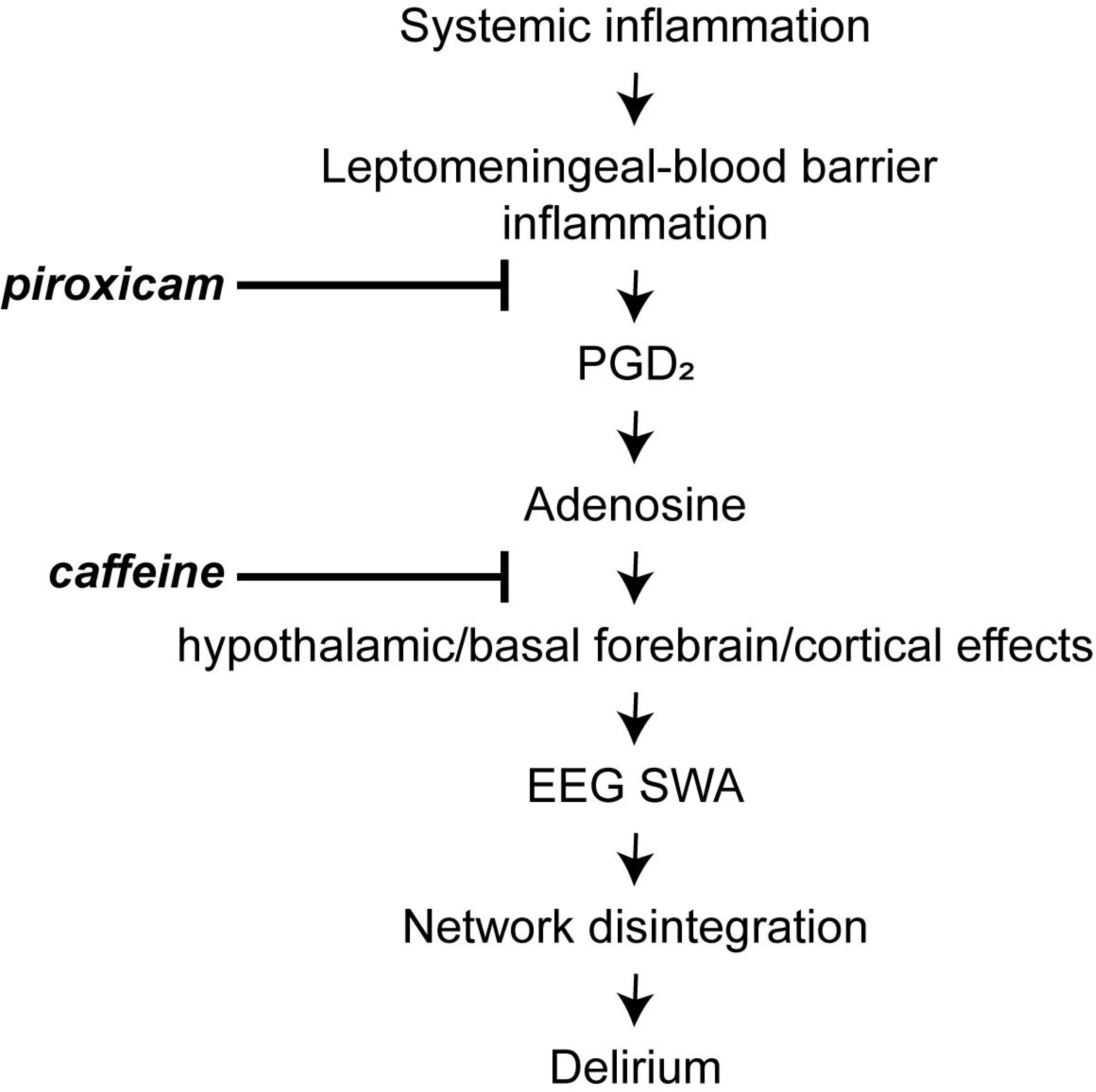
Working model of inflammation-driven SWA. Our working model of the generation of wakeful SWA relevant to delirium begins with systemic inflammation leading to Leptomeningeal-blood barrier inflammation, release of prostaglandin D_2_ (PGD_2_) and, via prostaglandin D receptor activation, release of adenosine. The enrichment of prostaglandin D receptors on the basal forebrain and hypothalamus suggests that adenosine then acts at the nearby arousal centres to promote sleep, and cortical SWA during wake. However, prostaglandin D synthase is diffusely expressed by the leptomeninges, and found in CSF, and so we cannot exclude additional direct cortical actions to induce SWA. Understanding whether the SWA reflects global or local slow waves seems a critical next step to determine whether delirium involves activation of sleep centres (likely inducing global slow waves that typically occur early in the night) or also local changes that support more restricted SWA (that may reflect local changes in adenosine related to PGD_2_ or metabolism).

In an exploratory analysis, we calculated power spectral density in overlapping 4-second windows and averaged the power spectra over epochs within lower or upper quartile movement ranges, as well as quiescence (Figure 2E *top*). As expected, under baseline conditions, the power spectra changed based on the magnitude of movement, with SWA suppressed during movement compared to quiescence, and in the highest quartile of activity compared to the lowest. Importantly, the same analysis applied to data recorded after administration of LPS did not show changes in the power spectra with increases in movement (Figure 2E *bottom*). Instead, following LPS administration, SWA was elevated even during movement, rendering power spectra during movement similar to those recorded during quiescence.

The analysis of Figure 2E suggests an effect of LPS on SWA during active wakefulness. However, to quantify this effect, we would want to compare SWA recorded during periods with identical movement profiles. To achieve this, we applied propensity score matching (Supplementary Figure 2) to compare SWA across comparable distributions of movement recorded during baseline and Peak LPS. LPS caused a steep increase in movement-matched SWA (Figure 3A), though there was no significant difference between LPS doses (Table 1G). SWA during quiescent periods was increased by LPS treatment (Figure 3B), though this effect was mainly driven by aged animals (discussed later) and no direct comparisons showed differences between the adult LPS-only groups (Table 1H).

### Cortical connectivity is unchanged in LPS after correcting for changes in activity

Under baseline conditions, we observed that alpha-band wPLI was positively correlated with movement (Supplementary Figure 6B; r^2=^0.435, p<0.05), suggesting that the decrease in wPLI observed in LPS could be due to the reduction in movement following injection of LPS. This is indeed what was observed when we applied propensity score matching to the relationship between wPLI and movement. To obtain accurate measures of wPLI, we expanded the time window for movement analysis to 20 seconds, slightly reducing the temporal resolution of changes in activity level. The movement-matched alpha-band wPLI was unaffected by LPS (Figure 3C; Table 1I), indicating the absence of changes in wPLI after accounting for LPS-induced decreases in movement. wPLI during quiescence was also unaffected by LPS treatment (Figure 3D; Table 1J).

### Neocortical cytokine levels correlate with changes in movement-matched SWA

We observed increases in cytokine levels following injection of LPS and systematic increases in SWA after accounting for the effect of LPS on movement. We next sought to determine if the magnitude of the increase in cytokine levels and the magnitude of the changes in brain activity were related for the measures that showed significant LPS effects. IL-6 levels correlated with increases in movement-matched SWA for LPS-only groups (Figure 4A; r^2=^0.662, p<0.00001), as did MCP-1 levels (Figure 4B; r^2^=0.541, p=0.0014).

### Piroxicam attenuates the EEG slowing in LPS without affecting brain IL-6 levels or acute decreases in movement

Piroxicam is a non-selective cyclooxygenase inhibitor previously shown to attenuate acute behavioral changes induced by LPS^3, 23^. We first determined that 10mg/kg piroxicam administration prior to Low LPS did not affect neocortical levels of IL-6 or MCP-1 compared to Low LPS alone (Supplemental Figure 7A-B; Table 1A), indicating any effect of piroxicam electrophysiologically or behaviorally would be downstream of the initial pro-inflammatory cytokine response, similar to prior reports.

Piroxicam blunted the LPS-induced increase of overall SWA (i.e. before accounting for movement; Figure 5A *red*; Table 1C) and gamma power (Supplementary Figure 4B; Table 1D) but did not decrease the impact of LPS on wPLI (Figure 5B; Table 1E). Piroxicam did not alter movement during the peak hours following LPS injection compared to Low LPS-only animals (Figure 5C). However, the difference in movement between baseline and the final recording hour (the fourth hour post-LPS, not included in ‘Peak LPS’) was significantly smaller in ‘PXM+Low LPS’ relative to Low LPS alone (likelihood ratio test: χ^2^(1)=16.625, p<0.0001), suggesting that piroxicam-treated animals may recover more quickly. Piroxicam attenuated the LPS-induced increase in movement-matched SWA (Figure 3A; Table 1G). As with Low LPS alone, piroxicam did not change SWA during quiescent periods (Figure 3B; Table 1H). Further, piroxicam animals showed smaller movement-matched SWA changes than would have been predicted from IL-6 (Figure 4A *red*; For PXM prediction residuals, one-sample t-test vs zero: t=-3.78, df=6, p=0.0092) or MCP-1 levels (Figure 4B *red*; for PXM prediction residuals, one-sample t-test vs zero: t=-3.54, df=6, p=0.012).

Given prior data showing that piroxicam blunts the prostaglandin response to LPS by inhibiting COX activity^3, 23^, and that prostaglandin D_2_ acts as a powerful somnogen via downstream effects on adenosine signaling^24, 25^, we further investigated the role of this pathway in LPS-induced SWA by testing the effect of caffeine citrate.

### Repeated injection of caffeine diminishes LPS-induced EEG changes

Caffeine promotes wakefulness through antagonism at the A_2A_ adenosine receptor^42, 43^, which is a downstream mediator of the somnogenic prostaglandin D_2_ that is known to affect cortical arousal^25, 44, 45^. When animals were administered LPS in combination with caffeine citrate, they showed lower overall SWA (Figure 5A *blue*), higher gamma power (Supplementary Figure 4B), increased alpha-band wPLI, and increased movement compared to animals administered LPS with saline (Figure 5B-C; Table 1C-F). Caffeine blunted the effects of LPS on the movement-matched SWA compared to animals treated with LPS plus saline (Figure 3A; Table 1G). Caffeine did not alter SWA during quiescence compared to saline-treated animals (Figure 3B; Table 1H). The effect of caffeine citrate treatment plus LPS on movement-matched wPLI relative to animals treated with saline plus LPS (Figure 3C) was strikingly similar to the effect of Vehicle versus Low LPS reported above (compare Figure 3C), though here the difference between the two was statistically significant (Table 1I). Caffeine pretreatment did not alter wPLI during quiescence relative to saline (Figure 3D; Table 1J).

### Aged animals exhibit exaggerated EEG slowing during quiescence

As delirium is most relevant in aged populations^46^, we repeated LPS experiments in aged mice and compared results to adult animals. As expected^47^, aged mice compared to adults showed higher neocortical IL-6 as well as higher MCP-1 protein levels in response to LPS treatment (Supplementary Figure 6A-B; Table 1A-B). Aged animals demonstrated an increased overall SWA response to LPS compared to adult animals (Figure 5A *green*; Table 1C), though this was not observed for gamma power (Supplementary Figure 4B; Table 1D). Decreases in wPLI were exaggerated in aged animals (Figure 5B; Table 1E). LPS-driven decreases in movement were not different between aged animals and adults (Figure 5C; Table 1F). Increases in movement-matched SWA due to LPS were not different from adult animals (Figure 3A; Table 1G). Instead, SWA during quiescence was greatly increased in aged compared to adult animals (Figure 3B; Table1H), and this was the only significant contrast in the model. As stated above, changes in both movement-matched and quiescent alpha wPLI following LPS treatment were not significant in models including aged animals (Figure 3C-D; Table 1I-J).

## Discussion

### Inflammation-induced changes in cortical activity

Inflammation causes acute changes in brain activity and connectivity in hippocampus, where changes in synaptic plasticity may underlie cognitive deficits observed during delirium^48, 49^. Less is known about the effects of inflammation on neocortical activity. Our results show some alignment with previous findings in rodents^22, 50^, where a much higher dose of LPS (1mg/kg) slowed hippocampal theta rhythms independently of changes in locomotion, though comparable effects were not observed in prefrontal cortex, suggesting the slowing is region-selective. The data presented here suggest a similar regional heterogeneity, as posterior increases in SWA were larger than anterior in High LPS animals, but this observation requires confirmation with further experiments. Posterior cortical SWA appears particularly important in delirium^14^. Previous studies have also described EEG slowing and decreases in alpha antero-posterior wPLI^50^, but did not account for effects on movement^40^ and also used a much higher dose of LPS (1mg/kg), making their results more likely to reflect somnolence rather than wakeful EEG activity^39^.

Our findings with aged animals indicate an exaggerated biochemical and electrophysiological response to inflammation, but since the differences between aged and adult animals were only observed in quiescence, those electrophysiological effects could be secondary to increased hypoactivity. Further work is needed to determine whether this rise in SWA in mice during quiescence reflects hypoactive wakefulness or sleep, but the data suggest that inflammation in aged animals induces a hypoactive phenotype that is particularly prevalent in elderly patients^46^.

### Inflammation-induced changes in cortical connectivity

The finding that alpha-band wPLI connectivity changes were attributable to changes in movement is potentially of clinical significance. The association of impaired alpha-band wPLI with delirium stems from work in postoperative patients where delirium is predominantly hypoactive^14, 29, 30^. In contrast, a study of delirium on emergence from anesthesia in young children, which is typically hyperactive, found increased alpha band connectivity during delirium^51^. In this context, reduced alpha-band wPLI may represent a specific marker of hypoactive delirium and this possibility should be tested in a cohort of patients including hypoactive and hyperactive delirium.

### Cytokine cascades contributing to inflammation

We acknowledge important differences in approaches to the study of inflammation in the mouse and human models. Notably we induced systemic inflammation in the mouse model using LPS, but focused on brain inflammation as a surrogate of neuroinflammatory hypotheses of delirium. In our recent work on delirium^14^ we studied plasma cytokines, a more distant surrogate of neuroinflammation. In the current study, we specified *a priori* that IL-6 would be the primary cytokine of interest due to its sensitivity to LPS^32^, long half-life^33^, and prior data from cerebrospinal fluid studies of delirium^52^. We complemented this with study of MCP-1 based on our recent paper^14^. Given redundancy in cytokine cascades, the next step is to understand local neuronal, immune, and circuit dynamics of these inflammatory stimuli. We suggest that inflammation drives EEG changes through induced release of prostaglandin D_2_ and subsequent effects on adenosine signaling, most likely in sleep and arousal centers in the hypothalamus and basal forebrain (Figure 6)^25^. Verification of the locus of adenosine’s actions, and investigation of possible direct actions of adenosine on cortical circuits, awaits further experiments. As non-steroidal anti-inflammatory drugs should be avoided in vulnerable elderly patients, future studies should focus on the therapeutic benefit of targeted manipulation of adenosine signaling in this mouse model and in a clinical setting.

### Translational relevance

Our focus has been on objective translational features of delirium that can be feasibly studied in the rodent. In contrast to studies of sepsis, where LPS is viewed as a suboptimal model^53^, LPS is commonly used in rodent studies of the mechanisms of delirium because it produces a profound and consistent proinflammatory response^3, 22, 49^. Our approach was to model how this inflammatory response affects EEG activity during movement to avoid the confound of sleep. As elevated SWA during wakefulness is a key criterion for delirium and is often associated with inattention, another key criterion, our model has a plausible association with delirium. The cognitive features that define delirium include impaired attention, arousal, executive function, orientation to the environment and memory as well as perceptual disturbances. Because inflammatory agents such as LPS affect motivation and motor function in animal models, investigating the cognitive correlates of inflammation behaviorally in mice is complicated^39^. Instead, we present an animal model of inflammation-related SWA during wakefulness; establishing this animal model opens opportunities for testing hypotheses about mechanisms and treatments for delirium and other inflammatory brain disorders. We consider this work to be complementary to work done with cognitive testing^3, 54^, which itself has limitations regarding the characterization of a complex human disorder in mice. Importantly, we recently showed that inflammation-driven SWA correlates with delirium severity in humans^14^, hence the ability to model this effect in mice and study mechanisms represents a major methodological advance. Furthermore, delirium, cognitive decline, and dementia are profound cognitive disorders associated with inflammation^2, 4, 5^ as well as changes in SWA^13, 55, 56^. Hence understanding the mechanisms whereby inflammation drives SWA may illuminate the key pathophysiological mechanisms of cognitive impairment in a variety of disorders.

### Future directions

The data presented here motivate future investigations into the mechanisms linking inflammation to changes in neural activity and connectivity. For example, previous studies have shown the acute behavioral effects of LPS are driven by peripheral IL-1B^49^; thus, it would be illuminating to measure plasma inflammatory markers in addition to measuring cortical cytokines. Future experiments should also identify specific receptor subtypes and other aspects of the signaling pathways involved in the link between neuroinflammation and changes in brain activity. For example, we tested caffeine, which is a non-specific adenosine receptor antagonist and has potential clinical application. Elucidating the roles of adenosine A_1_ versus A_2_ receptors would allow for more targeted drug development. In addition, future studies should include the effects of inflammation in mice modeling disorders associated with delirium, like dementia. More broadly, the model presented here opens opportunities for testing the roles of specific neuronal, glial, and immune cell types in the signaling cascade linking inflammation to drastic changes in brain function^55, 56^.

## Authors’ contributions

ZWS: Study design, data collection, data analysis, manuscript revision

ERJ: Study design, data collection, data analysis, writing first draft of paper

BMK: Study design, data analysis, manuscript revision

SMG: Data collection, data analysis, manuscript revision

CAM: Study design, manuscript revision

RDS: Study design, writing first draft of paper, manuscript revision

MIB: Study design, writing first draft of paper, manuscript revision

## Acknowledgements

The authors thank Elizabeth A. Townsend, Payge A. Barnard, Jens Mellby, Hazel Bastien, and Matthieu Darracq for technical assistance.

