## Supplementary for "Electrophysiological signatures of acute systemic lipopolysaccharide: potential implications for delirium science"

*Ziyad W Sultan, BS^a^

*Elizabeth R Jaeckel, BS^b^

Bryan M Krause, PhD^a^

Sean M Grady, BS^a^

Caitlin A Murphy, PhD^a^

Robert D Sanders, BSc (Hons) MBBS PhD DABA FRCA^c,d^

Matthew I Banks, PhD^a^

* these authors contributed equally

**Affiliations**

a – Department of Anesthesiology, School of Medicine and Public Health, University of Wisconsin–Madison

b – Department of Pharmacology, University of Michigan Medical School

c – Specialty of Anaesthetics, Faculty of Medicine and Health, University of Sydney

d – Department of Anaesthetics, Royal Prince Alfred Hospital, Camerpdown, New South Wales, Australia

**Corresponding Author**

Matthew I Banks, PhD

Professor

Department of Anesthesiology

4605 Medical Sciences Center

1300 University Avenue

Madison, WI 53706

608-261-1143

**Conflict of Interest**

The authors declare no competing financial interests

**Supplementary Methods**

**Animals**

All procedures with animals were approved by the University of Wisconsin-Madison Institutional Animal Care and Use Committee (IACUC) and in full accordance with Research Animal Resources and Compliance (RARC). Adult (2-8 months old; n=68) and aged (16-24 months old, n=14) c57Bl/6J mice were used in this study (Supplementary Table 1). [We note that within the broad age range in the ‘aged’ category, no significant relationship was observed between age and LPS-induced change in SWA (r^2^=0.20, p=0.11), IL-6 (r^2^=0.016, p=0.69), or MCP-1 (r^2^=0.0035, p=0.86).] Mice were obtained from Jackson Labs and maintained on a 12:12 reverse light-dark schedule (lights off at 09:00 AM) with *ad libitum* food and water for a minimum of 1 week prior to surgery and throughout the duration of the experiment.

**Electrode implantation**

Electrophysiological experiments were performed on a total of 72 mice (Supplementary Table 1). Mice were chronically implanted with four skull screw EEG electrodes using aseptic technique. Animals were anesthetized with isoflurane (1.5 – 2%) and placed on a heating pad to maintain temperature during surgery. After induction of anesthesia, the top of the head was shaved from the parietal skull bones to the frontal skull bones, an incision was made along the midline, and tissue was cleared to expose the skull bones. Craniotomies were drilled and stainless-steel screws were placed without piercing the dura mater bilaterally in the frontal (1.5 mm anterior to Bregma, 1.5 mm lateral to midline) and parietal (2.0 mm posterior to Bregma, 2.0 mm lateral to the midline) plates. Bilateral reference electrode screws were placed through the occipital plate and tied together to ground. EEG wires were silver (0.008” diameter, 0.011” diameter including insulation; A-M Systems, Sequim, WA) or stranded copper (0.012" diameter, 0.022" diameter including insulation; Cooner Wire, Chatsworth, CA). Wires were soldered to a 1-cm^2^ electrode interface board (EIB-16; Neuralynx, Bozeman, MT), which was fixed into place using dental cement (Fusio A3; Pentron; Orange, CA). Animals recovered in individual cages for at least 5 days prior to recording and were monitored for signs of distress and infection.

**Experimental paradigm**

Animal activity, resting-state EEG, and anterior-posterior functional connectivity were assayed 1 hour before and for 4 hours after LPS administration, after which animals were euthanized and their brains extracted and frozen for later cytokine ELISA (Supplementary Figure 1). The 4-hour duration of the post-injection recordings was chosen to be long enough to capture the peak effect of LPS while still allowing cytokine analysis afterward. Sterile 0.9% NaCl or LPS at doses of 12.5, 25, or 125 µg/kg were administered via intraperitoneal (IP) injection after the baseline hour. LPS was observed to exert its greatest effects on behavior and brain activity between 1 and 3 hours post-injection, and this time period was chosen for detailed analysis of treatment effects (see below). One animal from the 125 µg/kg LPS group was excluded from behavioral and EEG analysis due to poor signal quality but used for cytokine analyses. One or two days prior to each LPS experiment, animals were acclimated to experimental conditions using vehicle experiments with a saline injection instead of LPS.

Piroxicam EEG experiments lasted 6 hours, with IP injections of 10 mg/kg piroxicam (n=8, 4 female) and 25 µg/kg LPS administered after hours 1 and 2, respectively (Supplementary Figure 1, *red*). Brains were harvested 4 hours after LPS injection and snap-frozen until processing for cytokine analysis. Brains from un-implanted animals treated with 10 mg/kg piroxicam plus 25 µg/kg LPS (n=4, 2 female) and euthanized four hours post-LPS were also used in cytokine analysis.

Repeated IP injection of caffeine citrate was additionally tested as an intervention on the acute effects of LPS. Like LPS-only experiments, caffeine experiments followed a 5-hour timeline, except that injections of 30 mg/kg caffeine citrate (n=8, 4 female) or saline (n=10, 5 female) occurred at time of LPS (combined in syringe), followed by second and third injections of caffeine citrate alone at 1 and 2 hours post-LPS (Supplementary Figure 1, *blue*). Caffeine citrate was chosen based on its high aqueous solubility; 30 mg/kg was selected based on the relative masses of caffeine and citrate leading to an equivalent dose of 15 mg/kg caffeine, which has been shown to increase locomotion in mice^1^; repeated injections were chosen based on the short half-life of caffeine in mice^2^.

**Drug treatment**

All injections were prepared fresh daily, filtered using a 0.2µm filter (#66064-414; VWR International LLC, Pittsburg, PA), and administered IP at a volume of 5 ml/kg body weight. Frozen aliquots of ultrapure LPS from gram-negative *Escherichia coli* serotype 0111:B4 (#tlrl-3pelps; InvivoGen; San Diego, CA) in 0.01M phosphate-buffered saline (PBS; pH=7.4) were thawed, and the desired dose was diluted in 0.9% NaCl. Piroxicam (#P0847 Sigma-Aldrich; Atlanta GA) was dissolved in 0.2M Tris-HCl (pH=8) and administered at 10 mg/kg^3^. Caffeine citrate (#MP520481880; Fisher Scientific, Chicago, IL) was dissolved in sterile saline and administered at a dose of 30 mg/kg,

**EEG recordings**

All recordings were performed during the animals’ dark (i.e. active) period using equipment and software from Tucker-Davis Technologies (TDT; Alachua, FL). Individual animals were placed in a clear plastic beaker (6” diameter) within a dark sound attenuation chamber. A headstage (ZC16) with a flexible tether was attached to the electrode interface board and the animal was free to move about the chamber. Infrared cameras were trained on each cage at approximately a 45-degree angle to observe behavior. Initially, electrophysiological recordings were obtained with a RA16 preamplifier and RZ5 amplifier using BrainWare software (n=2; filtered at 2.2-12207 Hz, digitized at 24414 Hz). Later, recordings were obtained with a PZ5 preamplifier and RZ5D amplifier using either BrainWare (n=24; filtered at 0.4-12207 Hz, digitized at 24414 Hz) or Synapse software (n=45; filtered at 0.4-457 Hz, digitized at 1017 Hz), and stored for offline analyses.

**EEG analysis**

All data processing and analysis was performed using MATLAB software (Mathworks; Natick, MA) unless stated otherwise. EEG channels were excluded if visual inspection of the raw trace or spectra suggested the channel was compromised (n=21 channels of 76 channels in 19 animals, due to low signal amplitude, extreme sensitivity to movement artifact, or dominant electrical noise). Independent component analysis implemented in FieldTrip^4^ was used to remove apparent heart rate noise from one animal in the 125 µg/kg LPS group. All EEG data were downsampled to 200 Hz, then divided into 4-sec epochs (25% overlap) for power spectral density (PSD) estimation using Thompson’s multitaper method in FieldTrip. Noisy epochs were rejected from analysis if the standard deviation of the raw data in the epoch exceeded a set threshold. The threshold was set to the 95^th^ percentile of standard deviations within the index unless that value was below 0.2mV or exceeded 0.5mV. If the 95^th^ percentile was below 0.2mV the threshold was set to 0.2mV to prevent epochs from arbitrarily being rejected in clean data, and if the 95^th^ percentile exceeded 0.5mV the threshold was set to 0.5mV to ensure enough epochs were rejected in noisier data. Power was calculated in the following frequency bands: delta (i.e. SWA, 2-4 Hz), theta (4-12 Hz), alpha (13-20 Hz), beta (20-30 Hz) and gamma (30-80 Hz) and normalized by the mean total spectral power (2-80 Hz). The nonstandard low frequency cutoff of 2 Hz for SWA was used because two early recordings were obtained using an RA16 preamplifier with a high pass corner frequency hard-wired at 2.2 Hz.

Connectivity was assayed using the debiased weighted phase lag index (wPLI)^5^, a measure of phase synchrony, calculated in the alpha-band (13-20Hz) between viable anterior and posterior channel pairs on each hemisphere, and averaged across hemispheres. The alpha band was chosen as it is a standard connectivity metric used in human EEG studies^6,7^. For wPLI analysis, data were divided into 20-sec epochs (25% overlap), further divided into 0.5-sec trials, then a single wPLI value for each 20-sec epoch was calculated from the imaginary part of the cross spectral density obtained using the demodulated band transform^8^.

**Behavioral analysis**

LPS causes sickness behavior, evidenced as hunching, piloerection, and decreased locomotion^9^. Importantly, because the EEG power spectrum differs between inactive and active behavioral states^10^, to distinguish between changes in EEG parameters during wakefulness versus quiescence, epochs of recorded data were associated with a movement magnitude according to simultaneously recorded activity level. To monitor activity levels, video (mean frame rate $\geq$ 6 Hz in BrainWare or 10 Hz in Synapse; 240x320 pixels) was recorded for the duration of electrophysiological recording and analyzed offline. The video was read into the MATLAB environment using mmread^11^ and preprocessed in the following steps: 1) conversion from RGB to grayscale (values between 0-255); 2) masking via a user-drawn ROI to limit analysis within the frame to areas in which the animal could move; 3) rescaling the luminance values between 0 and 1 based on the 1^st^ and 95^th^ percentile luminance across the video; and 4) binarization based on each pixel being above or below a pre-determined threshold, with darker pixels assumed to be associated with the mouse. A frame-by-frame difference signal was then calculated to obtain a movement signal, defined as the power of the difference signal. A 2-D smoothing filter (MATLAB function ‘imfilter’) was applied to the movement signal, followed by time-domain smoothing (MATLAB function ‘smooth’) across approximately 2 seconds of video. Finally, the signal was averaged across all pixels, producing a 1-D vector with the estimated movement power in each frame of the video. The movement and EEG signals were then aligned in time, and the movement signal averaged in each 4-sec or 20-sec epoch for band power and wPLI analysis, respectively.

Epochs with nonzero estimated activity were used to calculate electrophysiological parameters corresponding to active wakefulness. To ensure that drug-induced changes in activity level for epochs classified as active wakefulness did not influence measured electrophysiological parameters, distributions of movement signal magnitude were matched between baseline and treatment periods using propensity score matching (PSM; Ho et al., 2007). PSM was applied to the movement epochs using the “MatchIt” package in R^12^ (Supplementary Figure 2) prior to calculating mean band power and wPLI for each animal. Movement distributions were matched using a full matching method, discarding windows from both baseline and peak effect hours with replacement. Animals with poor matching, defined as a greater than 1% difference in the mean movement distributions at baseline compared to peak effect hours after applying PSM, were excluded from the analysis (n=7/71 animals, distributed as follows: n=1 each from Low LPS, Aged Low LPS, and PXM + Low LPS, and n=2 each from High LPS, and Saline + Low LPS; Supplementary Table 1). Epochs with zero estimated activity throughout were used to calculate electrophysiological parameters corresponding to ‘quiescence’.

**Cytokine ELISA**

Both un-implanted animals (n=10) and animals implanted with EEG electrodes (n=46) were used for brain cytokine quantification. At the experimental endpoint, animals were deeply anesthetized with isoflurane, decapitated, and brains were removed, snap-frozen, and stored at -80°C until further processing. Whole brains were thawed on ice and dissected in ice-cold phosphate-buffered saline (PBS). Samples were obtained separately from anterior and posterior cortex, either by direct dissection or using a stainless-steel tube to extract cylindrical tissue samples, then homogenized via sonication in ice cold standard radioimmunoprecipitation assay buffer (RIPA; #20-188, Sigma-Aldrich, Atlanta, GA) with combined protease and phosphatase inhibitor tablet (PIA32955, VWR International LLC, Pittsburg, PA). Homogenates were centrifuged at 4°C and 17000xg for 15 minutes and the supernatant collected and refrozen until determination of total protein. Total protein concentration in supernatants was quantified using the Pierce BCA protein assay kit (Thermo Fisher 23225) according to the manufacturer’s instructions. Absorbance was read at 570nm on a Spectra MR (Dynex Technologies; Chantilly, VA) plate reader, and protein concentrations were quantified using a four-parameter logistic regression (elisanalysis.com). Prior to shipment, 103/111 samples were diluted with 0.01M PBS such that total protein concentration was equal to 1 mg/ml prior to ELISA. For samples with less than 1 mg/ml protein (n=8, range 0.73-0.97 mg/ml), statistical analysis was completed with and without corrections for protein content; no qualitative differences were found with corrected versus uncorrected values, so we present uncorrected data. One sample from a 125 µg/kg animal were excluded from analysis, as no total protein concentration value was obtained. All samples were analyzed undiluted, in duplicate, and fit to a standard curve ranging from 0.64 pg/ml – 10,000 pg/ml. Samples below the threshold of detection were imputed to the square root of the maximum lowest quantity observed (LOQ).

**Supplementary Table 1**

| **Test** | **Group** | | | | | | | **Figure** |
| --- | --- | --- | --- | --- | --- | --- | --- | --- |
|  | **VEH** | **Low LPS** | **PXM+Low LPS** | **Aged Low LPS** | **High LPS** | **Saline+Low LPS** | **CAF+Low LPS** |  |
| **Age (weeks)** | 20 ± 3 | 19 ± 3 | 19 ± 3 | 91 ± 13 | 24 ± 9 | 16 ± 6 | 15 ± 7 |  |
| **ELISA** | 5 | 18 | 12 | 13 | 8 | 0 | 0 | Fig. 1A-B, Supp. Fig. 7A-B |
| **Band power _all_** | 7 | 17 | 8 | 14 | 7 | 10 | 8 | Fig. 2B, 5A, Supp. Fig. 4A-B |
| **Band power _movt (PSM)_** | 7 | 16 | 7 | 13 | 5 | 8 | 8 | Fig. 3A |
| **Band power _qui_** | 6 | 16 | 7 | 13 | 5 | 8 | 8 | Fig. 3B |
| **Movt** | 7 | 16 | 7 | 13 | 5 | 8 | 8 | Fig. 2D, 5C |
| **wPLI _all_** | 7 | 17 | 8 | 12 | 7 | 10 | 8 | Fig. 2D |
| **wPLI _movt (PSM)_** | 6 | 16 | 7 | 11 | 7 | 6 | 8 | Fig. 3C |
| **wPLI _qui_** | 6 | 16 | 7 | 11 | 5 | 8 | 8 | Fig. 3D |
| **ELISA vs Band power _movt (PSM)_** | 5 | 11 | 7 | 0 | 5 | 0 | 0 | Fig. 4A-B |
| **Total (n _female_)** | 7 (1) | 23 (2) | 12 (6) | 14 (0) | 8 (0) | 10 (5) | 8 (4) |  |

**Supplementary Table 1. Animal groups and list of assays.** Animal groups, age at LPS experiment, and the animal counts for each test are shown, along with the corresponding Figures. CAF, caffeine citrate; LPS, lipopolysaccharide; movt, movement; PSM, propensity score matching; PXM, piroxicam; qui, quiescent; VEH, vehicle; wPLI, weighted phase lag index.

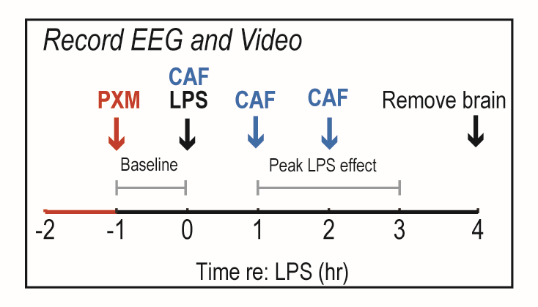

**Supplementary Fig 1**. **Experimental timeline and injection scheme.** Except for piroxicam and caffeine experiments (see below), EEG and video were recorded continuously for 1 hour (Baseline), followed by an IP injection of LPS. Recordings then continued for 4 hours, during which the behavioral effects of LPS were observed to peak (Peak LPS). After the recording period, mice were deeply anesthetized, decapitated, and brains were removed, snap-frozen, and stored at -80°C until ELISA. Baseline EEG and movement parameters were compared to the period 1-3 hours post-injection (Peak LPS). Piroxicam (PXM; *vermillion*) experiments included an extra hour at the start, followed by IP injection of 10 mg/kg piroxicam. Caffeine experiments (CAF; *blue*) followed a 5-hour timeline, but with injections of 30 mg/kg caffeine citrate or saline occurring at time of LPS (in syringe), and at 1 and 2 hours post-LPS. CAF, caffeine citrate; LPS, lipopolysaccharide; PXM, piroxicam.

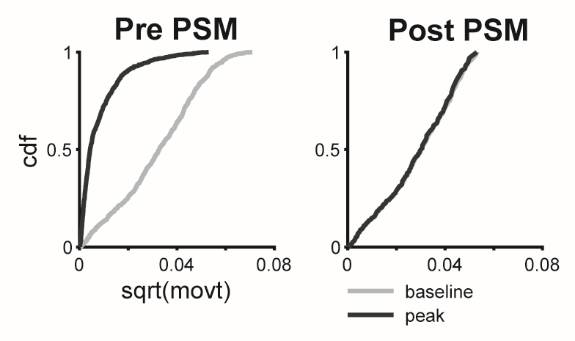

**Supplementary Fig 2. Movement distributions before and after propensity score matching.** Movement cumulative distribution functions at baseline (*gray*) or peak LPS (*black*) before and after applying propensity score matching (PSM).

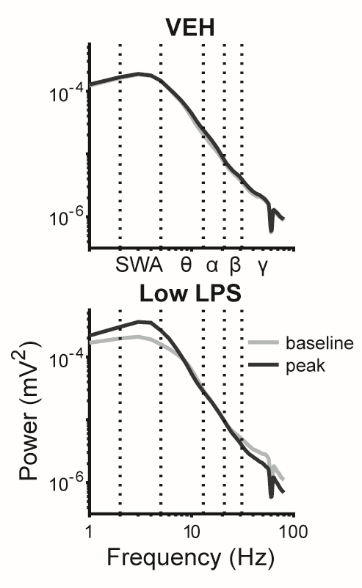

**Supplementary Fig 3. Power spectra at baseline and following LPS treatment.**

EEG power spectra visualized for the same animals and channels in 2A during the pre-injection hour (“baseline”, gray) and 1-3 hours post-injection (“peak”, black). Dotted lines indicate frequency band limits (slow wave activity [SWA], theta, alpha, beta, and gamma). LPS, lipopolysaccharide.

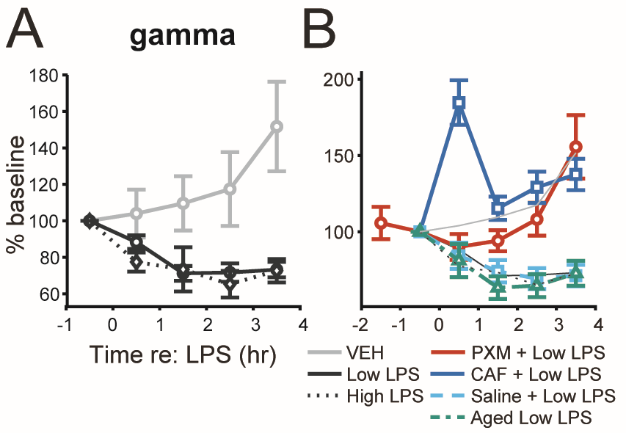

**Supplementary Fig 4. Normalized gamma power time series. A.** The time series of gamma-band (30-80Hz) power normalized to mean spectral power (2-80Hz) are shown for the different LPS doses. Symbols represent the mean percent change in gamma-band power from baseline across all animals at each LPS dose at each recording hour. Error bars represent ± SEM. **B.** Time series of normalized gamma-band power shown for additional animal groups. CAF, caffeine citrate; LPS, lipopolysaccharide; PXM, piroxicam; VEH, vehicle; wPLI.

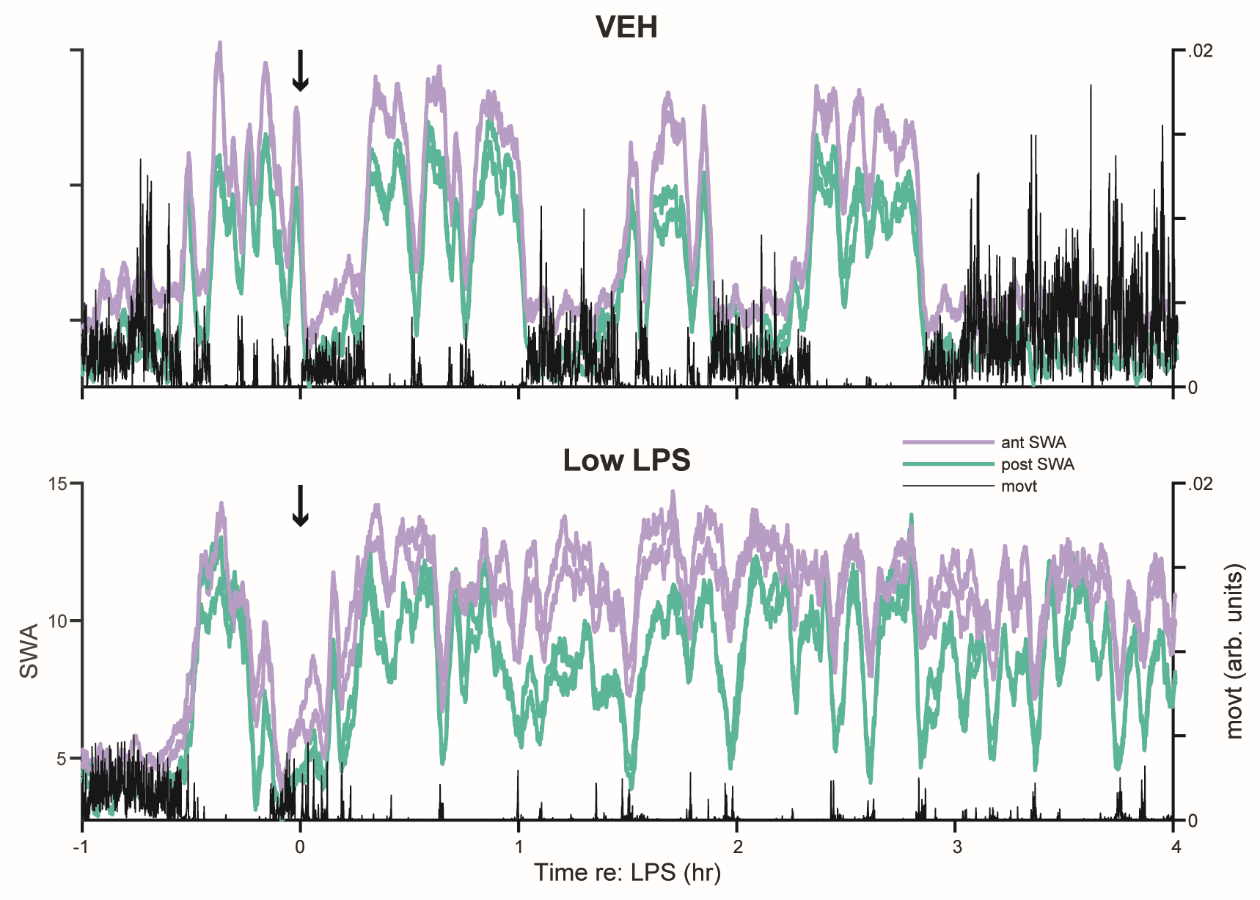

**Supplementary Fig 5. Behavioral impact of LPS and changes on SWA. A.** Time series of movement signals (*black*) and normalized SWA from anterior (*purple*) and posterior (*green*) EEG channels averaged over 4-second epochs shown for the same Vehicle (*top*) and Low LPS (*bottom*) mice as in Figure 2A and Supplementary Figure 3. Animal movement was estimated by analysis of infrared video recording, as described in the Supplementary Methods. LPS, lipopolysaccharide; SWA, slow-wave activity; VEH, vehicle.

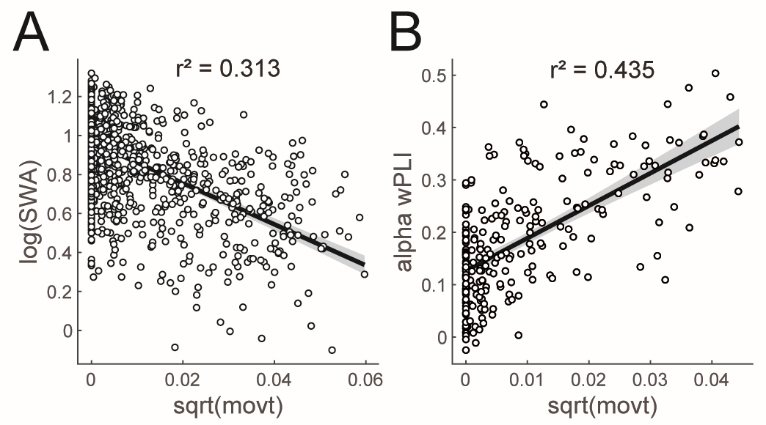

**Supplementary Fig 6. Movement correlations with EEG parameters. A.** Example correlation between log of EEG SWA and square root of movement (movt) from 1 hour of data at baseline. Each point represents the aligned SWA and movement signal from one 4-second epoch (n=1204). The line indicates the polynomial least-squares fit, and the shading indicates the 95% prediction interval of the regression line. **B.** Example correlation between alpha-band wPLI across anterior-posterior channel pairs and square root of movement from 1 hour of data at baseline. Each point represents the aligned alpha wPLI and movement signal from one 20-second epoch (n=240). SWA, slow-wave activity; wPLI, weighted phase lag index.

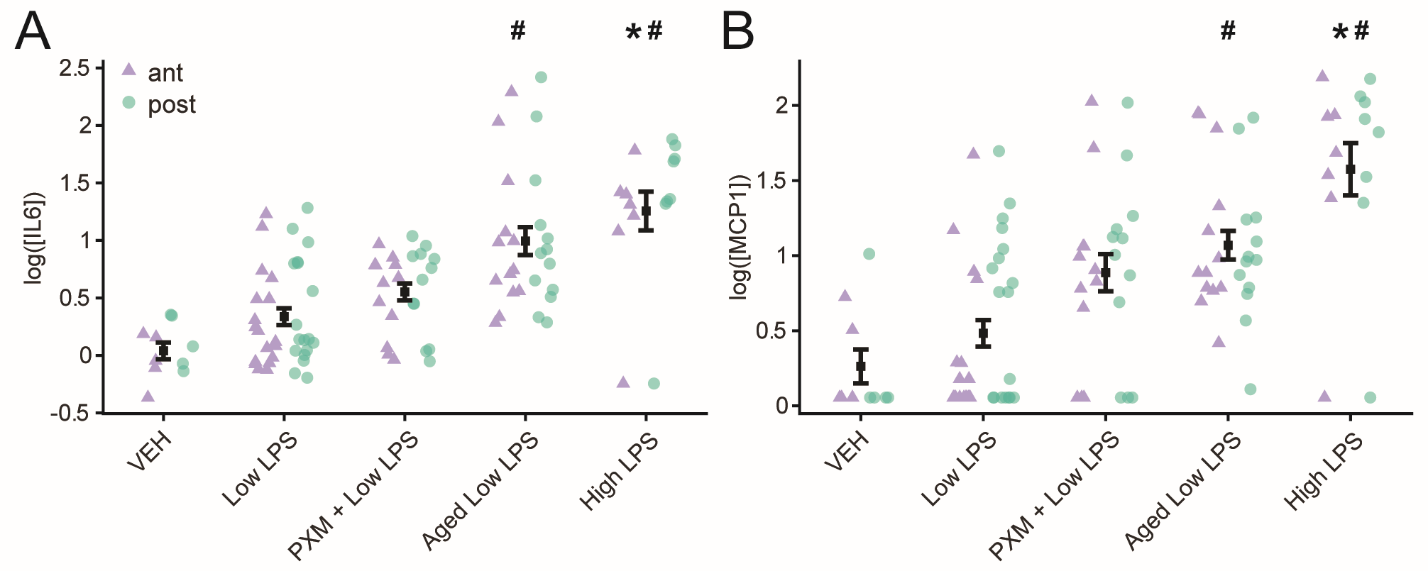

**Supplementary Fig 7. Proinflammatory cytokine levels in neocortex for all animal groups. A.** Neocortical IL-6 concentration shown for available animal groups (Vehicle, Low LPS, and High LPS are the same as in Fig 1; Caffeine + LPS and Saline + LPS animals were not included in the cytokine ELISAs). Each point represents log IL-6 concentration (pg/ml) from samples obtained bilaterally from anterior (ant; *purple triangles*) or posterior cortex (post; *green circles*). Overlaid symbols (*black*) represent the within-group mean across all samples. Error bars represent ± SEM. * indicates significant difference from Vehicle. # indicates significant difference from Low LPS. **B.** MCP-1 concentration shown for all available groups. IL-6, interleukin-6; LPS, lipopolysaccharide; MCP-1, Monocyte chemoattractant protein-1; SWA, slow-wave activity; VEH, vehicle.
